# Real-time accessible phylogenetics for every highly sampled virus

**DOI:** 10.64898/2026.09.10.750728

**Authors:** Angie Hinrichs, Lily Karim, Yatish Turakhia, Theo Sanderson, Russ Corbett-Detig

## Abstract

The scale of viral genome sequencing has outpaced the phylogenetic tools traditionally used to analyze it, as highlighted by the COVID-19 pandemic. We present viral_usher, a unified framework for scalable viral phylogenetics built on UShER. viral_usher is a containerized command-line tool that constructs mutation-annotated trees directly from public sequence repositories with minimal user input, building phylogenies of tens of thousands of genomes in minutes. Applying it across the International Nucleotide Sequence Database Collaboration, we assembled viral_usher_trees, a repository of 446 phylogenies spanning 163 well-sequenced viral species, rebuilt automatically monthly as new genomes are deposited. We extended Taxonium from a tree viewer into a web platform supporting in-browser phylogenetic placement and de novo tree construction, so that users can upload sequences and contextualize them within global phylogenies without local computational infrastructure. Because every tree is built by the same procedure, the repository enables comparative analyses across the breadth of viral diversity. We demonstrate the utility of this resource by asking what factors shape viral mutation spectra. We found that replication machinery, captured as Baltimore class, explains 46% of the variance across 162 viral genomes, while host taxon and envelope status together explain under 5%. These resources provide an extensible platform for real-time genomic epidemiology and for comparative evolutionary analysis across viral pathogens. Resources and code are freely available at https://taxonium.org/, https://github.com/lilymaryam/spectrum_analysis, https://github.com/AngieHinrichs/viral_usher_trees, and https://github.com/AngieHinrichs/viral_usher.

## Introduction

Pathogen genome sequencing has expanded dramatically as a consequence of the SARS-CoV-2 pandemic and widespread availability of genome sequencing technologies. Genomic epidemiology is becoming a critical element of basic biological discovery (Li, Grassly and Fraser, 2014), pathogen evolution analysis (Grenfell *et al*., 2004; Volz, Koelle and Bedford, 2013), infectious disease forecasting (Stockdale, Liu and Colijn, 2022) and public health response (Armstrong *et al*., 2019; Meredith *et al*., 2020). However, the scale and diversity of available viral genome data now pose substantial analytical challenges: datasets often comprise tens of thousands and up to millions of sequences, and exceed the capabilities of traditional phylogenetic inference methods, visualization tools, and computational workflows. These limitations highlight the need for scalable, automated, and broadly accessible frameworks for constructing, exploring, and maintaining large viral phylogenies.

During the SARS-CoV-2 pandemic, we developed the phylogenetic placement tool, UShER, to maintain a global phylogeny as it expanded to millions of sequences, with tens of thousands of genomes added daily (Turakhia *et al*., 2021; McBroome *et al*., 2021; Ye *et al*., 2022). In parallel, Taxonium was developed to enable interactive visualization of trees containing millions of sequences, and became essential for exploring these large phylogenies (Sanderson, 2022; Kramer, Sanderson and Corbett-Detig, 2023). Together, these tools enabled real-time phylogenetic analysis of pathogen genomes at an unprecedented scale, supporting genomic surveillance, lineage assignment, and outbreak investigation during the pandemic (Hinrichs *et al*., 2024; McBroome *et al*., 2024), with targeted applications to additional pathogens such as *Mycobacterium tuberculosis* (Karim *et al*., 2025). Despite these advances, constructing phylogenetic trees from public genome sequence data in the same manner for other viruses remains technically demanding, as it requires sequence retrieval, filtering, alignment, format conversion, and coordination of multiple software tools, which together create substantial barriers to entry and limit accessibility. Moreover, these steps would also have to be done routinely for maintenance of such phylogenies with database updates.

Here, we address both upstream and downstream challenges in large-scale viral phylogenetics. We present viral_usher, a containerized framework for automated retrieval, processing, and phylogenetic inference from public sequence repositories, producing mutation-annotated trees (MATs) with minimal user input. We further extend Taxonium to support real-time, in-browser phylogenetic placement via integration with UShER, letting users contextualize uploaded sequences within global phylogenies without local computational infrastructure. Building on these advances, we introduce viral_usher_trees, a public repository of regularly updated MATs spanning highly sampled viruses. Unlike tools such as Augur (Huddleston *et al*., 2021), which emphasizes maximum-likelihood inference at higher computational cost, our approach prioritizes scalability, accessibility, and automation, building phylogenies of thousands to tens of thousands of genomes in minutes.

Beyond improving accessibility, a unified framework for viral phylogenetics enables scientific questions that cannot be addressed when each pathogen is analyzed independently. Questions about what forces shape viral genome evolution are inherently comparative, and cannot be answered from any one virus. Yet existing resources are typically built independently for individual pathogens, making rigorous cross-virus analyses difficult. Because viral_usher_trees applies the same inference procedure uniformly across viruses, its mutation-annotated trees become directly comparable, permitting analyses that pool evidence across the breadth of viral diversity rather than treating each virus in isolation. We demonstrate this by asking what factors shape the evolution of viral mutation spectra, a question that has resisted resolution in part because no single study could assemble comparable trees across enough distinct lineages.

## Results and Discussion

### Viral_usher: A Containerized workflow for large-scale viral phylogenetic inference

Viral_usher automates the construction of MATs from public sequence data (Supplemental Figure S1). It is implemented as a Python command-line tool with two subcommands: init, which records all analysis choices to a structured configuration file, and build, which executes that configuration inside a Docker container. Separating configuration from execution keeps runs reproducible and lets init be scripted for batch use, while containerization removes the software-version and system-configuration discrepancies that are a frequent source of failure in multi-tool pipelines. The workflow retrieves sequences from NCBI, optionally adds user-provided genomes, filters them, aligns them to a reference with Nextclade (Aksamentov *et al*., 2021), and passes the resulting mutations to UShER, which builds and optimizes a MAT by maximum parsimony. Outputs include tabular metadata and a tree in UShER and Taxonium formats.

We evaluated the runtime performance of viral_usher across five viral datasets (Oropouche virus segment M, mumps, measles, RSV, and mpox) and three computing environments spanning typical user hardware to high-performance servers. Datasets under 2,000 sequences (Oropouche, mumps, measles) completed in one to two minutes on every platform tested, including sequence retrieval and preprocessing. Larger datasets exposed hardware differences: for RSV (>17,000 sequences) and mpox, a Linux server with 160 CPU cores and 1.5 TB RAM was up to two-fold faster than an Intel laptop and up to five-fold faster than an ARM laptop running under emulation (Supplemental Figure S2, Supplemental Table S1). Even the largest datasets ran to completion on a standard laptop, confirming that comprehensive phylogenetic inference is tractable without specialized computational infrastructure.

Viral_usher has significantly lower runtimes than Nextstrain’s Augur workflow, enabling the ongoing maintenance of up to date phylogenetic datasets for hundreds of viruses. On a 12-core ARM laptop, Augur required over 15 minutes for 1,231 measles genomes against under 1.5 minutes for viral_usher, and could not complete the RSV dataset at all, exhausting memory during multiple sequence alignment. On a high-performance Linux server, Augur took 30 to 35 minutes for measles and roughly 9 hours for RSV, compared to under 11 minutes for viral_usher (Supplemental Figure S2, Supplemental Table S1). Reference-based alignment and incremental parsimony placement avoid the computational bottlenecks of de novo alignment and maximum-likelihood inference, and this is what makes routine, repeated tree construction and augmentation from public data practical. That practicality is the basis for viral_usher_trees, a repository of viral MATs updated on a regular schedule as new genomes become available. Trees can be explored immediately in Taxonium or downloaded for local analysis with the UShER toolkit (McBroome *et al*., 2021, 2024; McBroome, Turakhia and Corbett-Detig, 2022; Sanderson, 2022; Turakhia *et al*., 2022).

### Viral_usher_trees: A monthly-updated repository of all highly sampled viral phylogenies

We applied viral_usher to publicly available genome sequences from the International Nucleotide Sequence Database Collaboration (INSDC; GenBank/ENA/DDBJ) to construct a repository of precomputed phylogenies in the form of MATs that is automatically updated monthly using GitHub Actions (https://github.com/features/actions). Starting from NCBI RefSeq reference genomes, we identified 672 candidate references across 196 viral species with substantial sequencing depth (≥1,000 publicly available genomes per species as of August 2025), excluding *Betacoronavirus pandemicum*, the species encompassing SARS-CoV-1 and SARS-CoV-2, given its exceptional scale and dedicated existing resources (Turakhia *et al*., 2021; McBroome *et al*., 2021). After requiring ≥80% genome coverage and acceptable ambiguity levels, and excluding datasets that retained fewer than 100 sequences passing filtering and aligning well to the RefSeq reference, we arrived at a final set of 446 MATs representing 163 distinct viral species, including 16 MATs for human-infecting viruses below the 1,000-genome threshold but above the threshold of 100 filtered and aligning sequences. These MATs are available through a GitHub repository (https://github.com/AngieHinrichs/viral_usher_trees), updated monthly using viral_usher’s incremental mode.

Building a unified resource across viruses requires choosing an appropriate root for each tree. RefSeq genomes provide a convenient coordinate system for mutation annotations, but they are typically modern, high-quality isolates rather than ancestral sequences, and rooting on them can distort interpretation for viruses with deep evolutionary histories. We evaluated three rerooting strategies: time-aware inference with TreeTime (Sagulenko, Puller and Neher, 2018), midpoint rooting on branch lengths (Farris, 1972), and outgroup rooting with external sequences (Felsenstein, 2004). TreeTime gives a principled estimate of the ancestral node but requires sampling dates. Midpoint rooting applies universally but assumes a roughly clock-like process (Hess and De Moraes Russo, 2007). Outgroup rooting can be robust but requires an appropriate external sequence and is sensitive to long-branch attraction (Bergsten, 2005). We applied the three methods in a fixed order of preference: TreeTime where sufficient sampling dates were available, midpoint rooting where they were not, and outgroup rooting where it placed the root more consistently with expected taxonomy than midpoint. This rooted 348 MATs with TreeTime, 94 by midpoint, and 4 by outgroup. After rerooting, we updated each tree’s reference sequence to match the inferred root while preserving the original RefSeq coordinate system. These references are not ancestral reconstructions but consistent coordinate frameworks that better reflect each tree’s topology, making the MATs more interpretable across diverse viruses and supporting comparative analysis in Taxonium.

The resulting MATs produced by viral_usher recover established lineage structure with high fidelity, demonstrating accurate local placement of sequences. For the 36 viruses carrying independent Nextclade clade assignments, we asked, for each of 698,141 labeled tips, whether its nearest neighbor by patristic distance belonged to the same lineage. Concordance was consistently high, with a median of 0.998 (range 0.934–1.000) and values at or above 0.99 for 31 of the 36 viruses. Because lineage labels were assigned by Nextclade against an external reference, independently of the tree topology we inferred, this agreement is not an artifact of internal consistency but reflects genuine recovery of lineage structure. Critically, the signal is not driven by clade-frequency imbalance, which could inflate concordance when one clade predominates. The median baseline, the concordance expected from clade frequencies alone, was only 0.34, giving a median excess over chance of 0.638. The strongest cases were the most demanding: RSV-A (45 clades, baseline 0.05) and dengue virus type 1 (69 clades, baseline 0.06) both reached concordance above 0.996, and concordance remained high in the largest MATs (0.998 for a H3N2 tree of ∼98,000 tips) and the most finely divided (0.980 across the 100 HIV-1 lineages). The speed of parsimony placement thus comes at little cost to the lineage relationships that surveillance and placement applications rely on.

**Figure 1.**
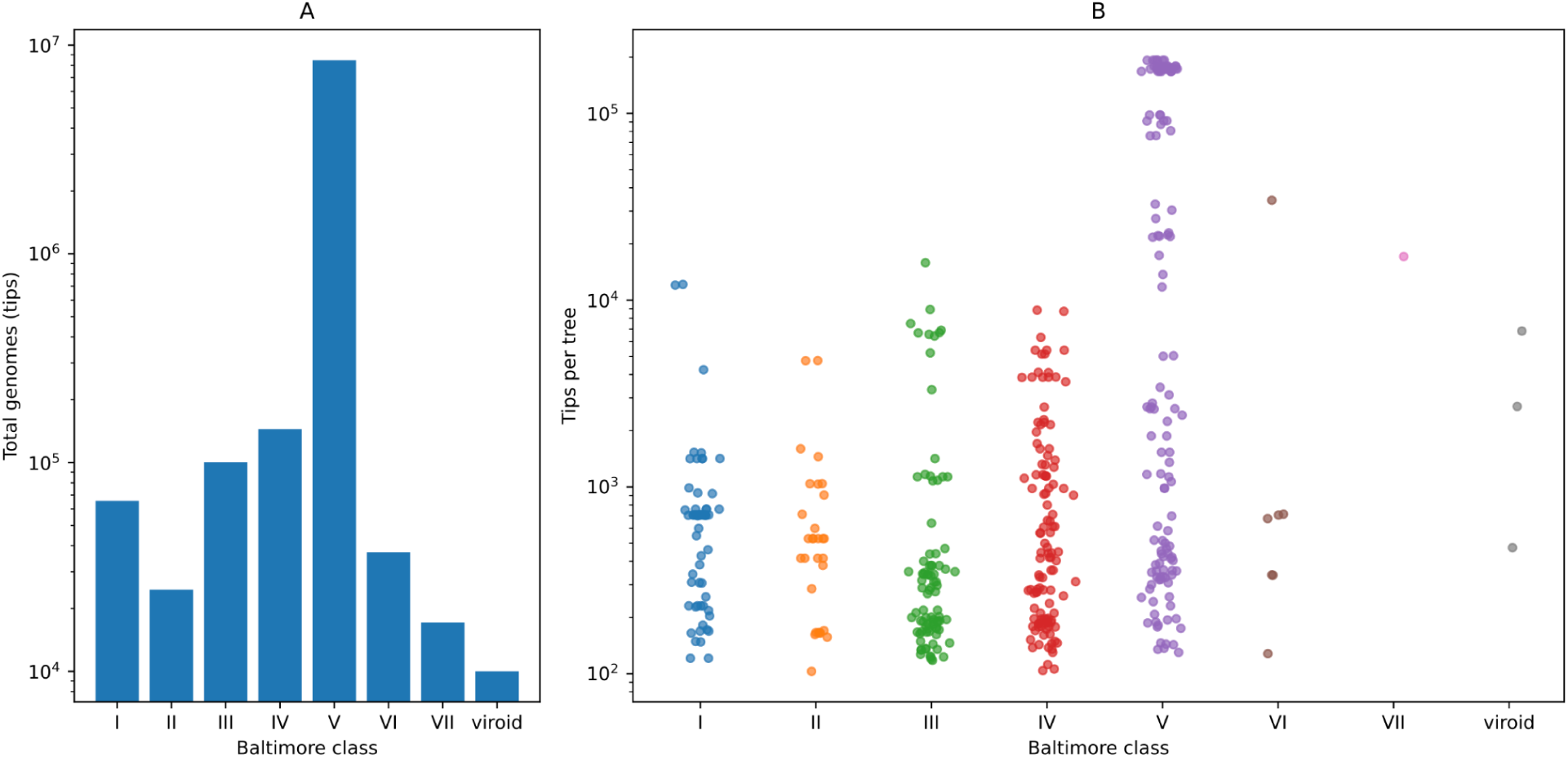
Phylogenetic resource composition by Baltimore class. Summary of the 446 viral phylogenies analyzed here, spanning 236 viral species and 8,863,599 total genomes (tips). Trees are grouped by the Baltimore class of the corresponding virus. The three viroid trees fall outside the Baltimore scheme and are shown separately. (*A*) Total genomes per class, summed across trees. (*B*) Tips per tree, with each point representing one tree and horizontal jitter added to reduce overplotting. Both panels use a log-scaled *y*-axis. (interactive version)

### Browser-based sample placement, tree construction, and scalable visualization

Even with improvements in automation, command-line workflows remain a barrier for many users, limiting both accessibility and the speed at which genomic data can be interpreted. To extend the reach of viral_usher and viral_usher_trees beyond computational specialists, we expanded Taxonium from a high-performance tree viewer into a full web-based platform for phylogenetic analysis. With this update, Taxonium retains its original capability to interactively visualize phylogenetic trees, including MATs, containing millions of sequences. Users are still able to navigate dense regions of the tree, zoom across scales, and query metadata in real time which are essential functions for exploring the increasingly large and complex datasets generated by global sequencing efforts. Our new features enable users to construct, explore, and analyze large viral phylogenies entirely within a web browser, eliminating the need for local software installation or specialized computational infrastructure.

To enable these features, we restructured the Taxonium architecture from a static single-page application into a dynamic, server-backed system. The frontend was migrated from React to Next.js, allowing seamless integration of client- and server-side operations, while a Python backend built with FastAPI orchestrates phylogenetic analyses. When users upload sequence data through the web interface, files are stored in cloud object storage and passed to backend services that automatically generate viral_usher configuration files and initiate analysis jobs on a Kubernetes cluster. These jobs execute the same containerized workflow used for command-line analyses, ensuring consistency between browser-based and local runs.

This system provides immediate feedback for running jobs in the form of logs streamed to the user interface, providing transparency into each stage of the workflow, from sequence retrieval and alignment through tree construction. Upon completion, resulting MATs are automatically integrated into the Taxonium viewer, where users can explore their data in the context of global phylogenies. Outputs are also made available for download, supporting downstream analysis in external tools.

In parallel, we developed a dedicated interface for interacting with the viral_usher_trees database. Users can browse and search across precomputed phylogenies spanning diverse viral species, rapidly identifying relevant datasets without needing to construct trees from scratch. For each tree, we provide integrated functionality for phylogenetic placement: users can upload new sequences and place them directly onto the existing global tree with a single action. This enables rapid contextualization of newly generated genomes, a critical requirement for real-time genomic epidemiology and outbreak investigation.

Because Taxonium makes viral_usher accessible through the web, it can easily be integrated with other tools as part of a broader bioinformatic ecosystem. We designed the platform such that Taxonium can accept, as URL-parameters, inputs that provide URLs to FASTA and metadata files. This allows an external tool to easily point users to Taxonium, preset to add a certain set of sequences to a specific viral_usher tree. We created an integration into the new Pathoplexus platform, which now allows users to use the Pathoplexus website (https://pathoplexus.org/) to select a set of sequences and then click a single button to reach the workflow to place them onto a tree, allowing the samples to be understood in phylogenetic context.

Together, these features fundamentally shift the accessibility of large-scale phylogenetic analysis. Tasks that previously required coordination of multiple tools, local compute resources, and technical expertise, such as building trees from raw data or placing sequences into global phylogenies, can now be performed interactively in minutes through a web browser. By tightly integrating automated tree construction, a continuously updated global database, and scalable visualization with in-browser analysis, Taxonium serves as the primary interface connecting users to the viral_usher toolkit, enabling both routine surveillance and exploratory, cross-virus evolutionary analyses.

**Figure 2:**
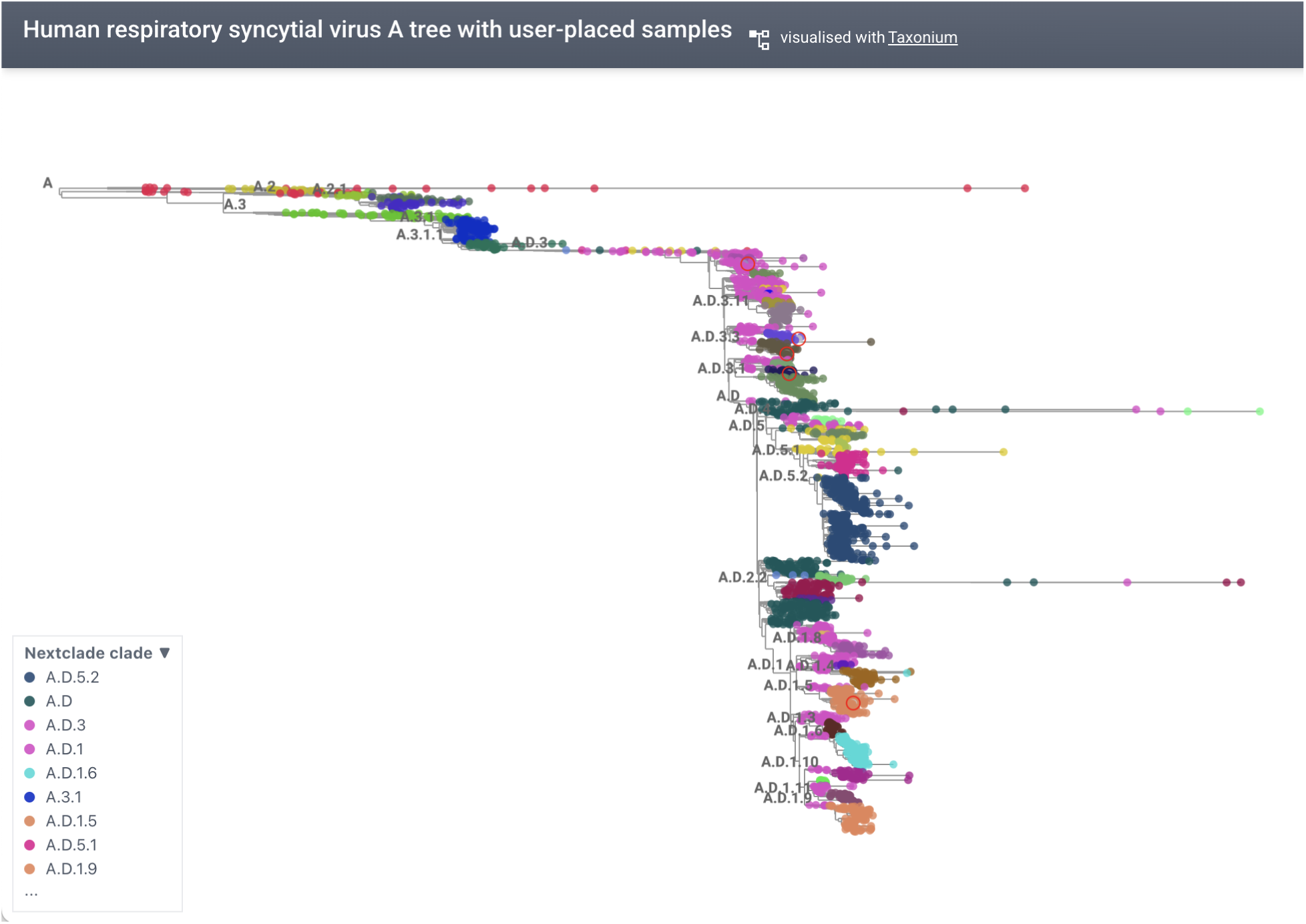
RSV-A tree with five user-provided sequences (red circles) added using Taxonium. An interactive view for user exploration is available from the Taxonium web platform (Interactive version).

### Application of viral_usher for Evolutionary Dynamics Inference

The mutation spectrum, *i.e.* the relative rates at which each nucleotide change occurs, shapes the supply of variation on which selection acts, making it a primary constraint on viral evolutionary dynamics. Its determinants nonetheless remain poorly resolved. One hypothesis holds that the viral environment, particularly host cell type, is the dominant driver, with host-supplied nucleotide pools, repair enzymes, and editing systems imprinting their signatures on the viral genome (Simmonds, 2020; De Maio *et al*., 2021; Ruis *et al*., 2023; Lamb *et al*., 2024). An alternative is that the replication and repair machinery encoded or recruited by the virus itself is the principal determinant (Corbett-Detig, 2025; Yang *et al*., 2025). This resource is uniquely positioned to address this concern.

The Baltimore viral classification scheme (Baltimore, 1971) offers a natural way to evaluate the relationship between replication mode and realized mutation spectrum. It sorts viruses into seven classes by the nature of their genome and the route they take to mRNA: double-stranded DNA (I), single-stranded DNA (II), double-stranded RNA (III), positive-sense (IV) and negative-sense (V) single-stranded RNA, and the reverse-transcribing RNA (VI) and DNA (VII) viruses. Because each route commits a virus to a particular replication apparatus, a host DNA polymerase, a virally encoded RdRp, a reverse transcriptase, and to particular intermediates that differ in how long the genome spends single-stranded and exposed, Baltimore class is a proxy for replication machinery rather than for host or environment. Our dataset of hundreds of distinct lineages spans all seven classes, so the two hypotheses can be compared directly.

Baltimore class is a major determinant of realized mutation spectra. Principal component analysis of mutation spectra from 162 viral lineages separates the classes cleanly along the first two components. Permutational multivariate analysis of variance (PERMANOVA) on Aitchison distances (Aitchison, 1982) shows that Baltimore class explains 46.2% of the variance in mutation spectrum (pseudo-F = 22.0, p = 10^-4^, 10,000 permutations), the single largest share attributable to any factor we examined. This reflects genuine differences in spectrum location rather than heterogeneous spread among classes: PERMDISP (permutational analysis of multivariate dispersions, Anderson, 2006) finds no difference in dispersion (F = 0.17, p = 0.99). By contrast, host taxon explains only 3.5% (p = 0.003) and envelope status 1.4% (p = 0.012) beyond Baltimore class. Nor is the Baltimore signal a proxy for host: fitting host first leaves Baltimore’s contribution essentially unchanged (46.3%), indicating that the two are effectively independent. Collectively this suggests that replication machinery is a major determinant of the viral mutation spectrum across the diverse lineages that we evaluated.

Even finer distinctions in replication mode may also leave their mark on the mutation spectra. Among ssDNA viruses (Group II), the G→T rate is strikingly bimodal. The high-rate mode (∼0.08–0.11) comprises exactly the CRESS-DNA viruses, *Circoviridae*, *Geminiviridae*, and *Nanoviridae*, while the low-rate mode (∼0.01–0.04) comprises the *Parvoviridae* together with the *Anelloviridae*. The split is complete, with no intermediate lineages. G→T is the canonical signature of guanine oxidation (8-oxo-G, Kamiya, 2004), and the two modes correspond to a difference in replication strategy. CRESS viruses replicate by rolling-circle replication, which leaves the genome single-stranded and unprotected for extended intervals, whereas parvoviruses replicate by rolling-hairpin replication from a linear template. Because these replication modes are largely conserved within major viral clades, phylogenetic relatedness could also contribute to the observed partitioning, making it difficult to fully disentangle shared ancestry from replication mechanisms. Nevertheless, the correspondence between replication strategy and a chemically interpretable mutational signature suggests that finer partitions of replication biology, beyond the Baltimore scheme itself, may explain variance currently residual to Baltimore class.

## Conclusion

Viral genome sequencing has outgrown the tools traditionally used to analyze it. Existing phylogenetic workflows are often fragmented, computationally intensive, and require substantial user expertise, limiting their accessibility and slowing the translation of data into insight. Here, we present a unified framework that replaces these constraints with a scalable, automated, and accessible system: viral_usher for rapid tree construction, viral_usher_trees for continuously updated global phylogenies, and Taxonium for interactive, browser-based analysis. Together, these components make it possible to move from raw genome sequences to interpretable phylogenetic context in minutes, without specialized infrastructure or advanced technical knowledge.

Our framework is best suited to questions where breadth, speed, and consistency matter more than the precision of any individual branch. Rapid contextualization of a newly sequenced genome, identification of mutations that have arisen independently across a phylogeny, monitoring the emergence and spread of a lineage during an outbreak, and comparison of evolutionary patterns across many viruses are all well served. viral_usher has already been incorporated in a metagenomics pipeline, demonstrating its utility in leveraging biobank-scale phylogenies to achieve haplotype-level resolution in clinical and environmental samples (Gangwar *et al*., 2026). For some applications, however, a maximum-likelihood or Bayesian inference may be preferable to parsimony-based placement using a reference-based alignment. For example, tests of selection at specific sites (Kosakovsky Pond and Frost, 2005; Yang, 2007) or sophisticated phylodynamic analyses (Baele *et al*., 2025) may warrant dedicated methods and expert curation. Importantly, our platform’s architecture also provides a natural foundation for future extensions, including ultra-scalable detection of recombinant lineages (*e.g*.. RIPPLES, Turakhia *et al*., 2022), maximum-likelihood optimization of parsimony-derived topologies (*e.g.,* MAPLE, De Maio *et al*., 2023) and scalable Bayesian phylodynamic inference (*e.g.* Delphy, Varilly *et al*., 2025). We view viral_usher and viral_usher_trees as complements to such analyses rather than replacements for them, and often as their starting point, supplying a curated sequence set and an initial topology on which additional phylogenomic analyses can be built.

This approach lowers the barrier to genomic epidemiology while improving speed, scale, and consistency across viruses. By standardizing data processing and enabling real-time interaction with global phylogenies, it supports both routine surveillance and rapid response in a way that existing approaches do not. As sequencing continues to expand, frameworks that prioritize automation, accessibility, and integration will be essential—and we expect this model to serve as a foundation for the next generation of large-scale evolutionary analysis.

## Data, Resource, and Code Availability

The viral_usher_trees repository includes files for trees in UShER and Taxonium formats and sample metadata, as well as all custom scripts described above, and is available at https://github.com/AngieHinrichs/viral_usher_trees. An interactive search page with links to download files or view in Taxonium is at https://angiehinrichs.github.io/viral_usher_trees/. https://taxonium.org/ also provides a search interface for visualizing trees as well as options to build a new tree or add your sequences to an existing tree. viral_usher is available from https://github.com/AngieHinrichs/viral_usher and can be installed using PyPi (pip install viral_usher) or Bioconda (conda install -c bioconda -c conda-forge viral_usher). Additional scripts used for analyses presented in this work are available from https://github.com/AngieHinrichs/viral_usher_trees/tree/main/scripts and https://github.com/AngieHinrichs/viral_usher_trees/tree/main/misc. Taxonium code is available at https://github.com/theosanderson/taxonium. Code for generating spectrum splits available at https://github.com/lilymaryam/spectrum_analysis with code for PCA analysis available at https://github.com/lilymaryam/spectrum_analysis/tree/main/viralUSherSpectra. Code for outgroup-based rerooting available at https://github.com/lilymaryam/outgroup_pipeline.

## Methods

### Viral_usher

#### Interactive species and reference genome selection

Viral_usher’s init subcommand begins by prompting the user for a viral species search term; the search term is sent in an NCBI Datasets (O’Leary *et al*., 2024) web query which returns a set of matching NCBI Taxonomy IDs and scientific names. The user selects a virus from the set, and then the Taxonomy ID is passed in an NCBI EUtils API query using BioPython’s Entrez module (Cock *et al*., 2009) to find RefSeqs associated with the Taxonomy ID. Matching RefSeqs are presented for the user to select one, which is used as the reference sequence for alignment and the root of the tree.

#### Data acquisition and preprocessing

Viral_usher’s build subcommand retrieves all viral genome sequences and metadata from INSDC (GenBank/ENA/DDBJ, *Nucleic Acids Research,* 2024) via NCBI EUtils API queries, NCBI’s datasets command line tool, and an NCBI Virus web query (https://www.ncbi.nlm.nih.gov/labs/virus/vssi/, Brister *et al*., 2015).

#### Sequence retrieval procedures (APIs, query logic)

Viral_usher’s build subcommand uses NCBI’s datasets command line tool to download the RefSeq sequence and gene annotations and to download all GenBank sequences for the Taxonomy ID when building a tree for the first time, or only new sequences when updating a previously built tree. It uses an NCBI Virus web query to download metadata for all GenBank sequences for the Taxonomy ID. When updating a previously built tree, it identifies new sequences by comparing the sequence accessions in the metadata to those already in the tree. The user can also provide a FASTA file containing additional sequences and a TSV file with metadata for the additional sequences to the init subcommand, to be used by the build subcommand.

#### Quality filtering

GenBank and user-provided sequences are filtered to a minimum length that is proportional to the length of the reference sequence (default 0.8 * reference length), and to a maximum proportion of Ns (default 0.25 * sequence length). If the same sequence name or ID is seen more than once then only the first instance is retained, with user-provided sequences processed before GenBank.

#### Alignment and variant calling

Sequences that pass filters are aligned to the reference using nextclade with default parameters, producing a multi-sequence alignment in reference coordinates that is piped to the faToVcf utility included in the usher bioconda package, which identifies substitution changes to the reference and outputs Variant Call Format (VCF). faToVcf’s output is compressed using gzip and used as an input to usher-sampled.

#### Phylogenetic inference

When building a tree for the first time, usher-sampled is run using the empty tree as the starting tree. In incremental update mode, the previously built tree is used. After usher-sampled has added the sequences from its VCF input, matOptimize is run to identify any branch moves that would improve the tree’s parsimony score. Then matUtils extract is run to filter sequences by configurable maximum number of private mutations (default 1000) and maximum branch length (default 10000 substitutions).

If the configuration file specifies a nextclade dataset for clade annotations then nextclade is run again to assign clades (most nextclade datasets include a reference that is not the same RefSeq used by viral_usher so clade assignment is run separately from alignment to reference), and matUtils annotate is used to label nodes of the tree corresponding to clade branches. Clade annotations are added to metadata from NCBI (and user-provided metadata if given).

If present in the metadata, then strain or isolate name, country and sample collection date are used to make a more descriptive name for each GenBank accession, and matUtils mask is used to rename all GenBank IDs in the final tree from the above steps (optimized.pb.gz), creating a separate tree file (viz.pb.gz) that uses the descriptive names. A metadata file indexed by the descriptive names and adding a numeric date column for spectrum coloring is generated and used as input to usher_to_taxonium (metadata.tsv.gz), along with gene annotations for the reference sequence, which writes the final output file for visualization with Taxonium (tree.jsonl.gz).

#### Workflow orchestration

All steps are orchestrated by a python script running in the docker image. If any step fails then the script exits with an error message.

Viral_usher has been tested on MacBook Pro laptops with Intel x86 and ARM CPUs, a linux server, GitHub Actions runner hosts, and running via kubernetes on a bare-metal x86_64 node.

### Viral_usher_trees repository

#### Reference genome selection

To identify reference genomes for viral_usher_trees, we began by searching the NCBI Virus web page (https://www.ncbi.nlm.nih.gov/labs/virus/vssi) for all RefSeqs associated with a viral species. As of August 2025, there were 18,740 viral RefSeqs. Using a custom script with EUtils queries, we determined the associated species-level Taxonomy ID for each RefSeq (many RefSeqs’ Taxonomy IDs are at a sub-species level which reduces the number of GenBank sequences that can be associated with the RefSeq). We then queried the number of GenBank sequences associated with the species-level Taxonomy ID, and retained the set of RefSeqs with at least 1,000 GenBank sequences, reducing the number of RefSeqs to 674. We excluded SARS-CoV-2 because a curated, daily-updated tree is already available (McBroome *et al*., 2021), and SARS-CoV-1 because it has the same species-level Taxonomy ID as SARS-CoV_2 which means that over 9 million GenBank sequences were found, with the vast majority of those being SARS-CoV-2. We generated viral_usher config files for the remaining 672 RefSeqs using a custom script, and built the trees using viral_usher on a Linux server. Some species have a high proportion of GenBank sequences that fail the filters on minimum length and/or number of ambiguous or ‘N’ bases, or are too diverged from the RefSeq to be aligned by nextclade with default parameters, resulting in fewer than 100 GenBank sequences being included in the tree. We discarded those RefSeqs and trees, resulting in a set of 432 RefSeq-based trees of at least 100 GenBank sequences. We later discarded two trees because their RefSeqs were redundant with other RefSeqs for the same species and subtype, and added 16 trees for human-infecting viruses that had fewer than 1,000 GenBank sequences but at least 100 sequences included in the tree, resulting in a final set of 446 trees.

#### Automated monthly updates

The GitHub Actions platform is used to perform an automated monthly update of all trees in the viral_usher_trees repository, orchestrated by a workflow file and driven by a tree metadata file. Multiple batches of tree update processes are run in parallel, with batch size constrained by the maximum run time of a GitHub actions job (6 hours), with the number of parallel running jobs limited by the number of queries that can be sent within a short time to NCBI (about 5 in the absence of an API key). For each tree in viral_usher_trees, viral_usher uses the existing optimized.pb.gz file as the starting tree, and attempts to add all GenBank sequences listed in metadata that are not already in the tree. On successful completion of viral_usher, output files are committed to the repository. Inevitably, a few of the 446 update jobs fail due to repeated NCBI query failures, so failures are tolerated but recorded in a file. After all jobs have completed, the workflow tags a new release.

Addition of a new tree is still a manual process: the tree metadata file must be updated, a new subdirectory and config file must be created, an initial run of viral_usher must be performed locally, and the rerooting process must be repeated.

#### Rerooting procedures

We ran TreeTime only on trees for which the metadata included sample collection dates for at least 80% of sequences in the tree. A custom script extracted sample collection dates, determined whether there were enough well-formatted dates to run TreeTime, used matUtils extract to get a Newick representation of optimized.pb.gz with branch lengths in units of substitutions, used TreeSwift (Moshiri, 2020) to scale branch lengths to TreeTime’s expected units of substitutions per site, ran treetime clock, and extracted the ID of the node with the oldest estimated date to use as the new root. TreeTime completed successfully for 340 of the 432 trees in the set at the time of this comparison, and for 16 of the 16 trees that were added later.

The midpoint of each tree was computed by a custom script that used the simple algorithm of finding the internal node most distant from root, finding the internal node most distant from that node, and computing the midpoint of the path between the two nodes, taking branch lengths into account. Since midpoint is a function of a tree with no dependence on metadata or any external data, it was computed for all trees.

We built a Snakemake pipeline to identify a potential outgroup for each viral tree (available from https://github.com/lilymaryam/outgroup_pipeline/). The pipeline searched for candidate outgroups by BLASTing each reference genome against a database of RefSeq viral genomes and selecting the highest-scoring hit outside the target taxon (excluded by NCBI taxonomy ID). RefSeq was searched first to prioritize curated reference sequences; when it yielded no hit, BLAST was rerun against a database of viral GenBank sequences. For each virus with a candidate outgroup, the corresponding sequence was downloaded and aligned to the reference with MAFFT. The alignment was converted to a VCF and placed onto the existing viral tree using the UShER subtool usher-sampled, without re-inferring the rest of the phylogeny. The tree was then rerooted to the parent node of the outgroup using matUtils, and the outgroup was pruned to yield the candidate rooted phylogeny for comparison against the other rerooting methods.

#### Reference/root sequence selection

We visually evaluated the rerooted trees produced by all methods that ran successfully for each tree and selected the rerooting that looked best by subjective criteria, with a bias toward treetime when treetime ran successfully. The viral_usher config file for each tree was updated to use the rerooted tree and modified reference sequence (having the same nucleotide coordinates as the original RefSeq, but modified base values to match the new root node).

#### Lineage Consistency Analysis

To assess whether the trees recover established lineage structure, we tested, for each tree carrying independent clade assignments, whether tips of the same clade are placed as nearest relatives. Clade labels were taken from the nextclade_clade field of each tree’s metadata, which is assigned by Nextclade against its own reference tree, independently of the UShER topology being evaluated. Trees without clade annotations were excluded, as was Sudan ebolavirus, whose metadata contained a single informative clade. This left 36 viruses.

For each tree, we loaded the optimized mutation-annotated tree (optimized.pb.gz) with the Big Tree Explorer (McBroome, Turakhia and Corbett-Detig, 2022) and defined the patristic distance between two tips as the number of substitutions along the path connecting them, taking each branch length to be its count of mutations. For every labeled tip, we identified its nearest labeled tip by patristic distance and recorded whether the two shared a clade; tips lacking a clade assignment were excluded both as focal tips and as candidate neighbors. Nearest neighbors were computed exactly in linear time by a two-pass dynamic program over the tree — a postorder pass recording, for each node, the nearest labeled tip within its subtree, and a preorder pass propagating the nearest labeled tip outside it — rather than by all-against-all comparison, which is intractable for trees of this size. Ties in distance, including the genetically identical sequences common in densely sampled trees, were broken arbitrarily.

We summarized each tree by its concordance, the fraction of labeled tips whose nearest labeled neighbor shared its clade. As a null expectation, we computed a baseline equal to the probability of two randomly selected sequences having the same clade (Simpson index), the concordance expected if a tip’s neighbor were drawn according to clade frequencies alone. The excess of concordance over this baseline measures agreement attributable to tree structure rather than clade imbalance.

### An Expanded Taxonium Web Platform

#### Frontend

For this project we re-implemented the Taxonium website in Next.js to enable server-side functionality. This package is built using Typescript and includes the modularised Taxonium viewer as a React component. We added a new section to the front page for exploring the viral_usher_trees dataset, a new facility for searching the now much larger dataset of trees, and new /build and /place pages. The /build page permits the construction of a new tree de novo, while the /place page allows users to place their own sequence onto an existing tree from viral_usher_trees.

#### Backend

We built a new backend service to handle phylogenetic placement, implemented in Python using FastAPI. This contains an endpoint for exploring the viral_usher_trees dataset, to permit searching for specific trees. It also supplies the endpoint called to build a tree. This endpoint accepts various parameters from the web form, which can include file uploads. It uploads the files in question to an S3 based storage, generates a viral_usher config file with the desired parameters, and then spawns a viral_usher job in the Kubernetes cluster with the generated config file. Another endpoint permits retrieval of logs as the job runs.

The kubernetes jobs each comprise two containers, a viral_usher container using the docker image described above, and a small sidecar that watches for completion of the job and then uploads the resulting output files. Source code is available from https://github.com/theosanderson/taxonium.

### Mutation spectrum comparative analyses

#### Dataset processing

To analyze mutation spectra for each virus in viral_usher_trees, we developed a Snakemake pipeline which extracted the rerooted viral phylogeny in Mutation Annotated Tree format (MAT) and adapted an existing mutation spectrum analysis codebase intended to analyze phylogenetic datasets in MAT format (Corbett-Detig, 2025). The pipeline follows approximately the same preprocessing structure as in Corbett-Detig, with a few notable changes for generality described below (available at https://github.com/lilymaryam/spectrum_analysis/).

The first preprocessing step prunes internal nodes that exceed a certain mutation:tip ratio to prevent outliers and possible aberrant data from confounding the algorithm. Instead of choosing a fixed mutation:tip threshold of 3 as in Corbett-Detig, we allowed the algorithm to calculate a threshold of three standard deviations above the mean mutation:tip ratio, which accommodates the smaller and sparser viral trees compared to the global SARS-CoV-2 phylogeny of the original analysis.

After pruning these long branches, we followed the Corbett-Detig approach for masking individual positions whose relative mutation rates increased sharply at a particular node in the phylogeny. We again modified the original approach, because parameter cutoffs appropriate for the SARS-CoV-2 global tree would not generalize across 432 smaller, heterogeneous datasets. We modified the position-masking algorithm to calculate both a minimum number of mutations required for a position to be considered for masking and a **χ**^2^ threshold for masking, set by a Bonferroni correction over the number of sites and nodes tested (α = 0.05).

We then calculated the mutation spectra of the entire trees (without looking for splits in mutation spectra as done in Corbett-Detig) by entering a very large **χ**^2^ threshold that inhibited identification of subsequent splits. This produced a mutation spectrum computed at the root node of each tree, which was used for downstream analysis. To minimize the impact of pseudoreplication, we further removed any dataset where the sample set overlapped another by 5% or more.

#### Mononucleotide mutation spectra

We estimated mononucleotide mutation spectra from the mutation-annotated trees in viral_usher_trees using the spectrumSplits tool (Corbett-Detig, 2025). For each reference genome, this yielded counts of each of the twelve possible mononucleotide substitutions (A→C, A→G, A→T, C→A, and so on), summed across the tree.

Raw substitution proportions confound the underlying mutation process with genome base composition, because a given substitution type X→Y can only occur at sites currently occupied by base X. A genome rich in A will therefore appear enriched for A→N substitutions regardless of its mutational process. To remove this dependence, we counted the number of unambiguous A, C, G, and T bases in each RefSeq genome (excluding N and other ambiguity codes) and divided the proportion of each substitution type by the genomic frequency of its source base. The twelve resulting per-site rates were rescaled to sum to one, giving opportunity-normalized spectra that are directly comparable across genomes of differing base composition.

Segmented viruses are represented in viral_usher_trees by one tree per segment. To avoid pseudoreplication, spectra from segments of the same virus were collapsed into a single per-virus spectrum by averaging the twelve normalized rates, weighting each segment by its total number of substitutions. Because each segment spectrum sums to one, the weighted average does as well. This reduced 273 segment-level spectra to 162 per-virus spectra. Each virus was assigned to a Baltimore class, and annotated for envelope status and primary host taxon, according to family-level assignment. The three viroids in the dataset (hop stunt, peach latent mosaic, and potato spindle tuber) have no Baltimore class and were retained as a separate category.

#### Principal component analysis

Principal component analysis was performed on the 162 × 12 matrix of opportunity-normalized substitution rates. Features were mean-centered but not scaled to unit variance, since all twelve are expressed in the same units, and components were obtained by singular value decomposition. Variance explained by each component was computed as its squared singular value divided by the total. Convex hulls around each Baltimore class in the PC1–PC2 plane in Figure 3 are drawn for visualization only and do not represent confidence regions.

**Figure 3.**
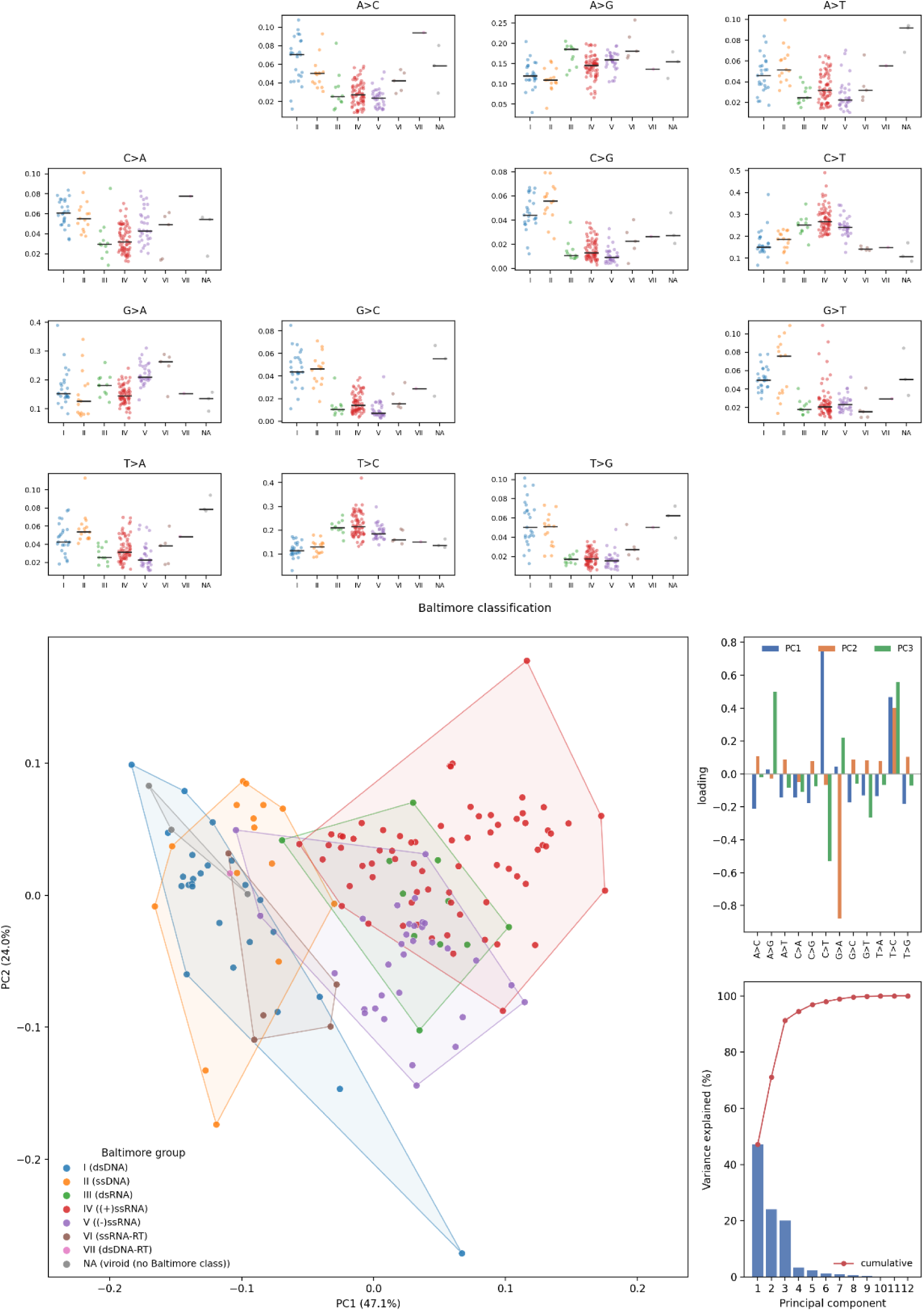
Principal component analysis of mutation spectra across viruses, grouped by Baltimore class. (Main panel) PCA of the resulting per-virus spectra; each point is one virus, colored by Baltimore group, with shaded convex hulls delimiting the extent of each group. Axes are annotated with the percentage of total variance explained by PC1 and PC2. (Right, top) PC1–PC3 loadings for each of the twelve mutation types, indicating which substitutions drive separation along each component. (Right, bottom) Scree plot showing per-component and cumulative variance explained. (Top grid) Per-virus rates for each mutation type, arrayed as a 4×4 matrix of from-base (rows) by to-base (columns) panels with the diagonal omitted, points jittered within Baltimore group and horizontal bars marking group medians. (interactive version)

#### Permanova

To partition variance in the mutation spectrum among predictors, we used distance-based permutational multivariate analysis of variance (PERMANOVA). Because the twelve normalized rates are compositional, constrained to sum to one, Euclidean distance is not appropriate; we instead used Aitchison distance, computed as the Euclidean distance between centered log-ratio (CLR) transformed spectra. Zeros were handled by adding a pseudocount of half the smallest positive value in the matrix prior to the CLR transform.

We fit sequential (Type I) models of the form spectrum ∼ Baltimore class + envelope status + host taxon, in which each term’s R² reflects the variance it explains beyond the terms preceding it. Pseudo-F statistics were assessed against 10,000 permutations of sample labels. Because Type I sums of squares depend on term order, we also fit the model with host taxon entered first, to test whether the Baltimore signal was confounded with host. Baltimore class VII was represented by a single virus (hepatitis B) and was excluded from the test, leaving 161 viruses in seven groups.

PERMANOVA is sensitive to heterogeneity of multivariate dispersion: groups differing in spread rather than in centroid location can produce a significant pseudo-F. We therefore accompanied each test with PERMDISP, comparing the mean distance of each virus to its group centroid in Centered Log Ratio (CLR) space, with significance assessed by 10,000 permutations.

## Acknowledgements

The authors thank the many researchers, laboratories, public health agencies, and sequencing initiatives that generated and deposited viral genome sequences in GenBank. This work relies on the collective contribution of the global scientific community in making viral genomic data openly available. We also thank the National Center for Biotechnology Information (NCBI) and GenBank for maintaining these resources and providing access to the sequence data used in this study. AI was used to enhance the readability of portions of the manuscript, and to prototype or extend portions of the code.

## Funding Acknowledgements

This study was funded in part by CDC 75D30124C20302 to RC-D. TS is supported by 306920/Z/23/Z from Wellcome.

---

**Supplementary Figure S1:**
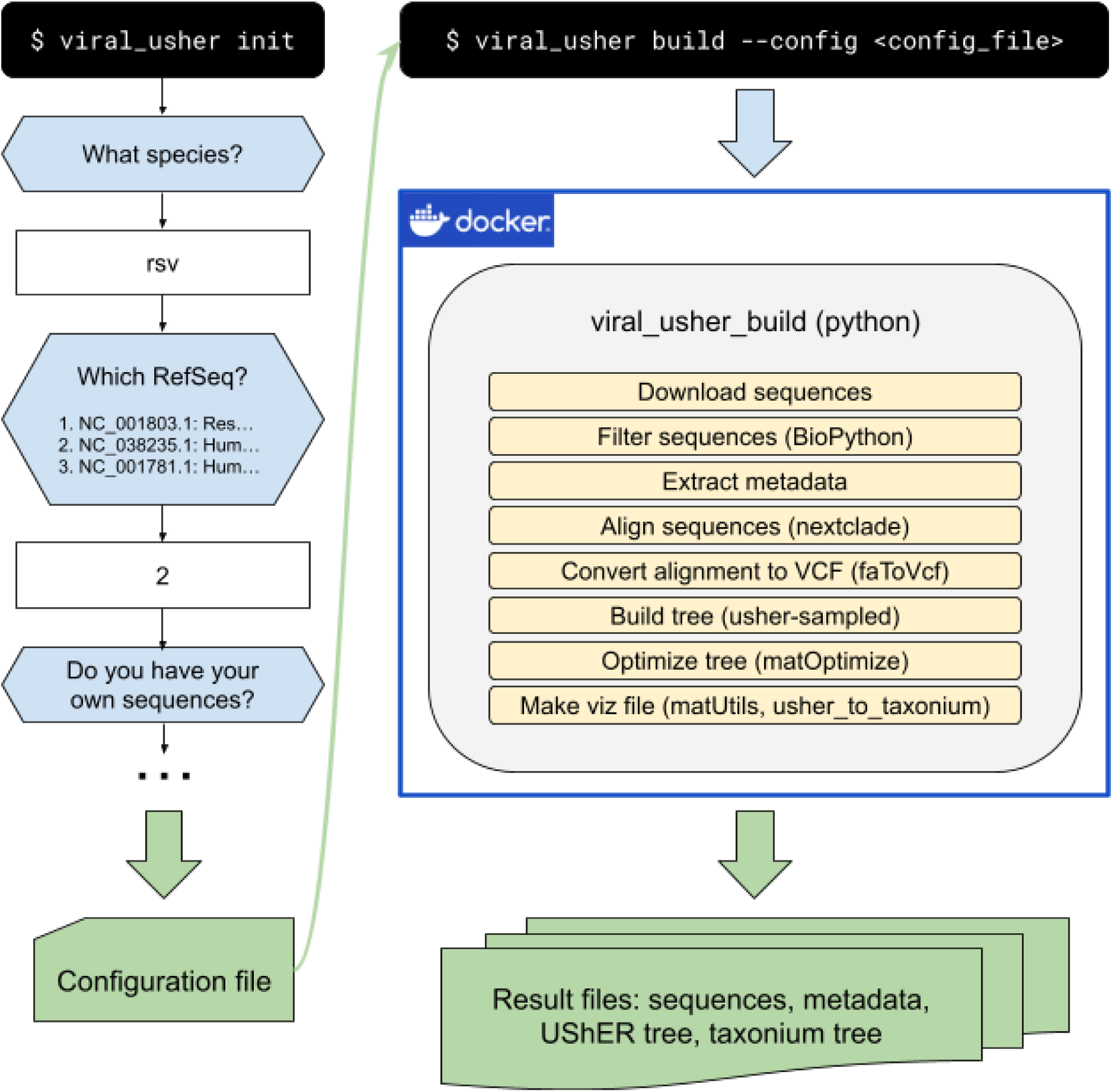
viral_usher is run in two stages. First, the init subcommand interactively queries the user and generates a configuration file recording the user’s input. Then the build subcommand follows the configuration file’s specification to download and process sequences and metadata and build the tree in a Docker container.

**Supplementary Figure S2:**
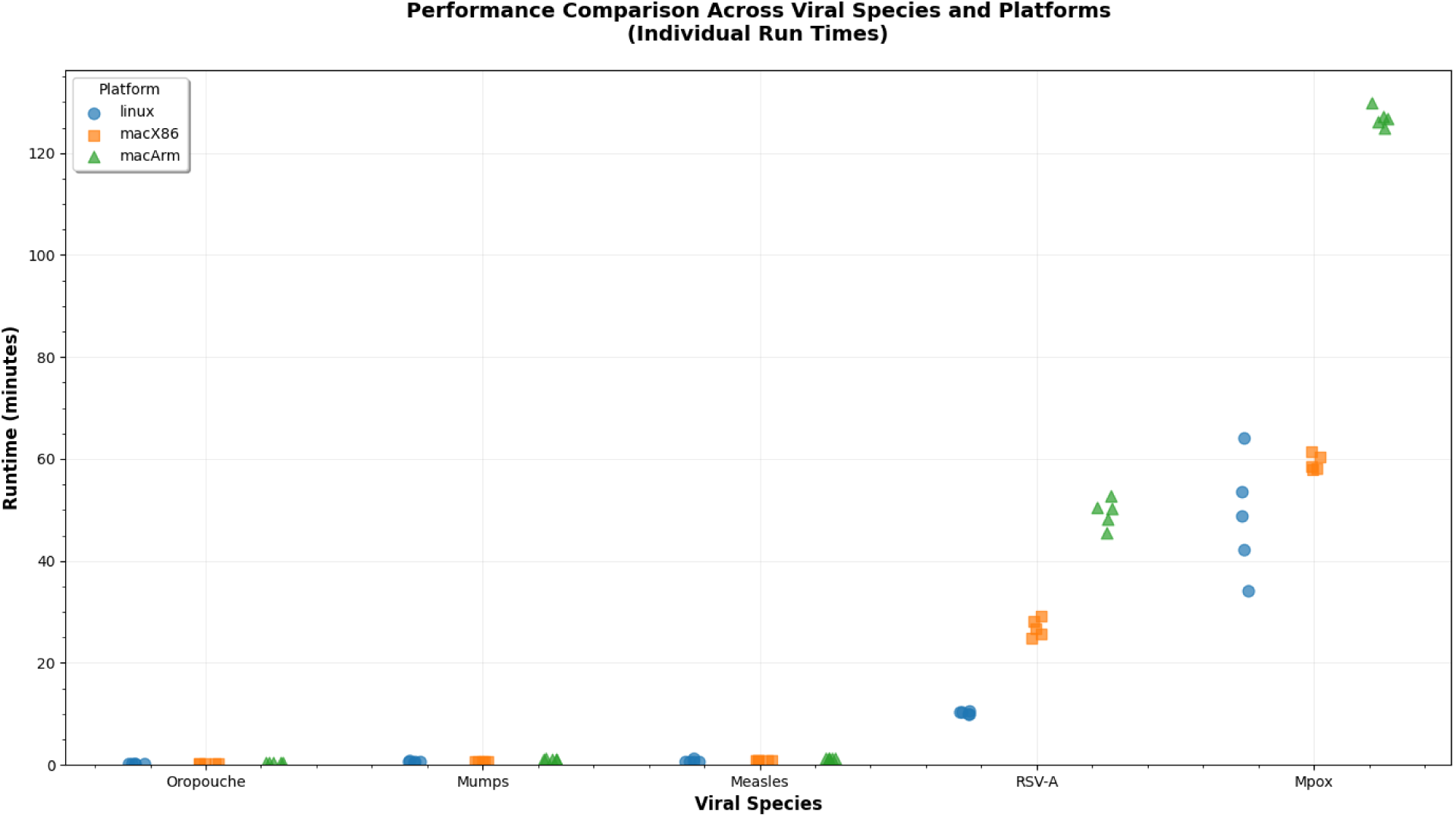
Run times of five trials on three platforms (shared linux server, MacBook Pro with Intel CPU, MacBook Pro with ARM) for Oropouche virus, Mumps virus, Measles virus, RSV-A and Mpox virus. Run times are very short for the first three viruses with small genome size and 1200 or fewer full-length sequences available at the time of testing, but differences between platforms become more apparent with RSV (17,000 sequences) or Mpox (genome size almost 200,000 bases).

**Supplementary Table S1:** Run times (min, median, max) and peak RAM usage as measured by docker stats on three platforms for five viruses of varying size and number of available near-full-length sequences. Only the M segment of Oropouche was used to build a tree, reducing the number of available sequences significantly. The subtype A reference was used for RSV and the clade I reference was used for Mpox. Longer sequences and larger numbers of sequences aligned to reference increase the run time. Performance on a newer Mac with a 12-core ARM CPU is worse than on an older Mac with a 6-core Intel CPU, presumably because the linux/x86 platform must be emulated on the ARM platform.

| Virus | RefSeq length | GenBank genome count | Filtered count | Aligned count | Has Nextclade dataset | Linux run time; peak RAM | Mac x86 run time; peak RAM | Mac ARM run time; peak RAM |
| --- | --- | --- | --- | --- | --- | --- | --- | --- |
| Oropouche | 4385 (M) | 2629 | 1597 | 800 (M) | no | 0.29m, 0.30m, 0.30m; 209.3M | 0.24m, 0.26m, 0.28m; 116M | 0.50m, 0.51m, 0.52m; 71.1M |
| Mumps | 15384 | 14584 | 1334 | 1334 | no | 0.56m, 0.57m, 0.79m; 707.4M | 0.61m, 0.63m, 0.67m; 144.4M | 1.07m, 1.12m, 1.19m; 304.7M |
| Measles | 15894 | 24693 | 1231 | 1231 | yes | 0.63m, 0.64m, 1.25m; 869M | 0.81m, 0.82m, 0.84m; 246.2M | 1.28m, 1.36m, 1.38m; 356.5M |
| RSV | 15222<br>(A) | 58971 | 17630 | 17627 | yes | 9.92m,<br>10.37m,<br>10.65m;<br>6.005G | 24.82m,<br>26.63m,<br>29.19m;<br>3.161G | 45.60m,<br>50.32m,<br>52.69m;<br>3.423G |
| Mpox | 196858<br>(1) | 10584 | 9941 | 9935 | yes | 34.19m,<br>48.91m,<br>64.21m;<br>28.52G | 58.00m,<br>58.53m,<br>61.40m;<br>6.001G | 124.80m,<br>126.86m,<br>129.83m;<br>6.966G |

